# Genomic risk score offers predictive performance comparable to clinical risk factors for ischaemic stroke

**DOI:** 10.1101/689935

**Authors:** Gad Abraham, Rainer Malik, Ekaterina Yonova-Doing, Agus Salim, Tingting Wang, John Danesh, Adam Butterworth, Joanna Howson, Michael Inouye, Martin Dichgans

## Abstract

Recent genome-wide association studies in stroke have enabled the generation of genomic risk scores (GRS) but the predictive power of these GRS has been modest in comparison to established stroke risk factors. Here, using a meta-scoring approach, we developed a metaGRS for ischaemic stroke (IS) and analysed this score in the UK Biobank (n=395,393; 3075 IS events by age 75). The metaGRS hazard ratio for IS (1.26, 95% CI 1.22-1.31 per standard deviation increase of the score) doubled that of previous GRS, enabling the identification of a subset of individuals at monogenic levels of risk: individuals in the top 0.25% of metaGRS had a three-fold increased risk of IS. The metaGRS was similarly or more predictive when compared to established risk factors, such as family history, blood pressure, body mass index and smoking status. For participants within accepted guideline levels for established stroke risk factors, we found substantial variation in incident stroke rates across genomic risk backgrounds. We further estimated combinations of reductions needed in modifiable risk factors for individuals with different levels of genomic risk and suggest that, for individuals with high metaGRS, achieving currently recommended risk factor levels may be insufficient to mitigate risk.

## Introduction

Stroke is a leading cause of death worldwide and the leading cause of permanent disability^1,2^. About 80% of stroke cases are of ischaemic origin^3^. The risk of ischaemic stroke is determined by a complex interplay of genetic and environmental factors partly acting through modifiable risk factors such as hypertension and diabetes. Roughly thirty-five genomic loci have been robustly associated with stroke^4–7^, and many more genetic associations have been reported for stroke-related risk factors^8–14^, e.g., over 1,000 loci have been associated with blood pressure (BP)^11,15–19^ and >100 with atrial fibrillation (AF)^10,20^. These data are now beginning to be harnessed to aid risk prediction.

Recent work has highlighted the potential of genomic risk scores (GRS) for risk prediction of common diseases^21–24^. Genomic risk prediction has a notable advantage over established risk factors as it could be used to infer risk of disease from birth, thus allowing the initiation of preventive strategies before conventional risk factors manifest and their discriminative capacity begins to emerge.

For stroke, a recent 90-SNP GRS derived from the MEGASTROKE GWAS meta-analysis^4^ showed that genetic and lifestyle factors are independently associated with incident stroke^24^, and that even among individuals with high GRS, lifestyle factors had a large impact on risk, implying that risk could be reduced in those with high genetic predisposition for stroke. However, in contrast to GRSs for other cardiovascular diseases like coronary artery disease (CAD)^21–23^, the predictive power of previous GRS for stroke has been limited^25–27^, likely because of limited genetic data for stroke and the well-known heterogeneity of the stroke phenotype^4,7^. Recent analytical advances have enabled more powerful GRS construction, such as those leveraging multiple sets of GWAS summary statistics^21,28^, potentially allowing for power and heterogeneity limitations to be overcome. Specifically, for CAD, an approach, where multiple GRSs are combined into one metaGRS, was found to improve risk prediction over any one of the individual CAD GRS^21^. Such an approach may be widened to provide substantively improved genomic prediction of stroke.

Here, we extended the metaGRS strategy to predict ischaemic stroke by incorporating GWAS summary statistics for stroke and its etiological subtypes ischaemic stroke (IS) as along with GWAS summary statistics for those of 19 other risk factors and co-morbidities of ischaemic stroke (IS). This new IS metaGRS was validated and compared to previously published GRS using UK Biobank^29,30^. We next compared the predictive capacity of the IS metaGRS to established non-genetic risk factors for IS. Finally, we assessed the additional information provided by the metaGRS in combination with current guidelines for the treatment of established IS risk factors and created joint models which predict absolute risk of incident IS.

## Results

### Derivation of a metaGRS for ischaemic stroke

To create the GRSs we randomly split the UK Biobank (UKB) British white dataset (n=407,388) into a derivation (n=11,995) and validation set (n=395,393; **Methods**, **Figure 1**, **Table 1**). In order to increase statistical power in the derivation phase, we enriched the derivation set (n=11,995) with ischaemic stroke events (n=888, 7.4%). A schematic of the overall study design is given in **Figure 1**.

**Figure 1:**
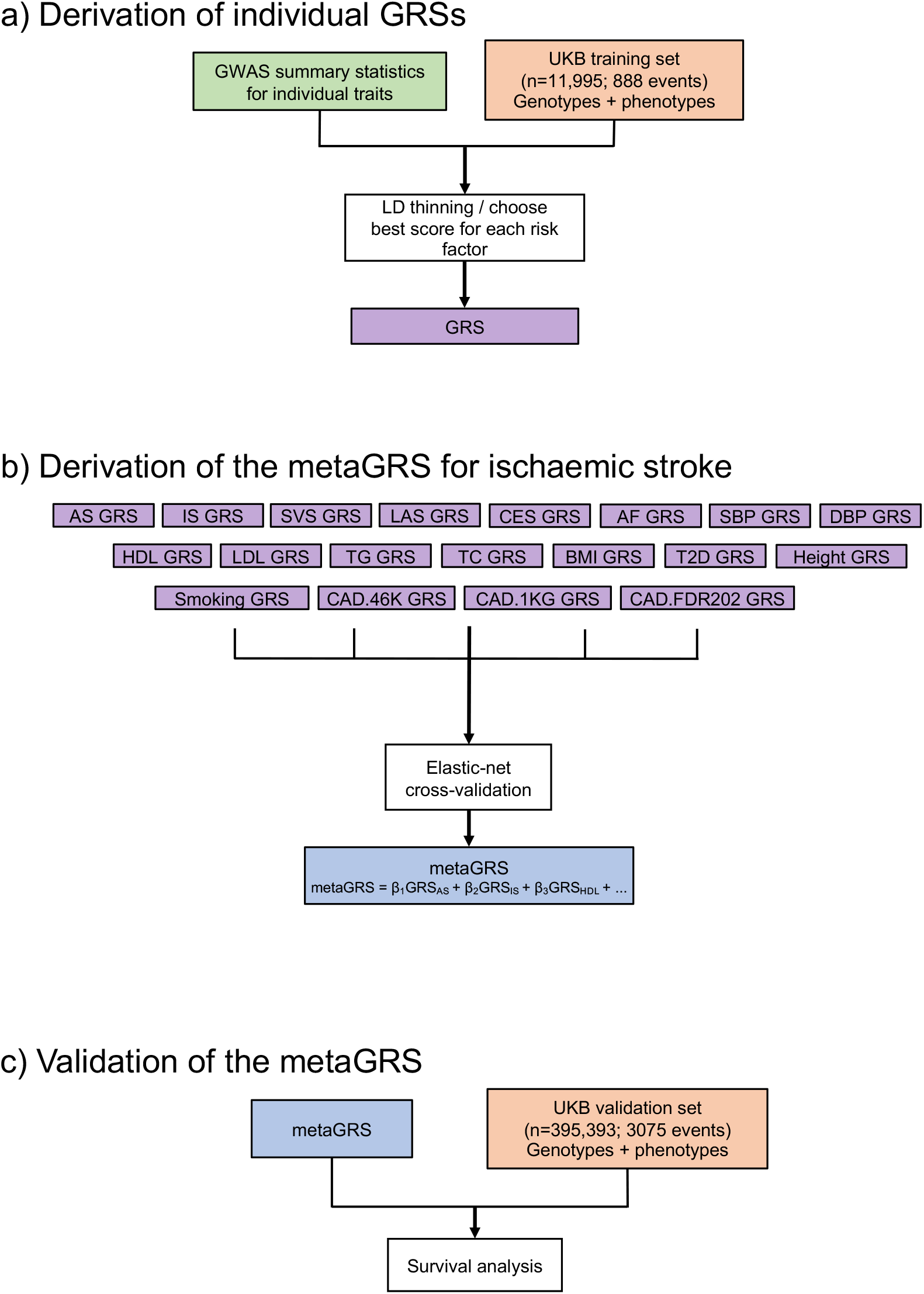
Study design. (a) Individual GRSs were derived in the UK Biobank training set (n=11,995) using GWAS summary statistics for individual traits. (b) The metaGRS for ischaemic stroke was then derived by integrating individual GRSs using elastic-net cross-validation. (c) Validation of the metaGRS for ischaemic stroke was performed in the UK Biobank validation set (n=395,393). Abbreviations: UKB, UK Biobank; GWAS, genome-wide association study; GRS, genomic risk score.

**Table 1:** Study characteristics of the UK Biobank validation dataset. Shown are characteristics obtained at the first UK Biobank assessment.

| Baseline characteristic | UK Biobank<br>N=395,393 | Male<br>N=180,653 (45.7%) | Female<br>N=214,740 (54.3%) |
| --- | --- | --- | --- |
| Age, years [mean (sd)] | 56.9 (8.0) | 57.1 (8.1) | 56.7 (7.9) |
| Current smoker, N (%) | 39,804 (10.0%) | 21,261 (11.8%) | 18,543 (8.6%) |
| Systolic blood pressure, mm Hg<br>[mean (sd)] (adjusted for BP<br>medication) | 143.3 (21.7) | 146.9 (20.4) | 140.2 (22.2) |
| Diabetes diagnosed by doctor, N<br>(%) | 18,675 (4.7%) | 11,449 (6.3%) | 7,226 (3.4%) |
| Hypertension, N (%) | 211,069 (53.4%) | 110,540 (61.2%) | 100,529 (46.8%) |
| Family history of stroke, 1<sup>st</sup><br>degree relative, N (%) | 104,831 (26.5%) | 45,569 (25.2%) | 59,262 (27.6%) |
| High cholesterol, N (%) | 53,141 (13.4%) | 30,670 (17.0%) | 22,471 (10.5%) |
| Prevalent stroke events, N (%) | AS: 4543 (1.1%) | AS: 2679 (1.5%) | AS: 1864 (0.9%) |
| AS: any stroke | IS: 1152 (0.3%) | IS: 787 (0.4%) | IS: 365 (0.2%) |
| IS: ischaemic stroke |  |  |  |
| Before age 75 |  |  |  |
| Incident stroke events, N (%) | AS: 2607 (0.7%) | AS: 1531 (0.8%) | AS: 1076 (0.5%) |
| Before age 75 | IS: 1923 (0.5%) | IS: 1207 (0.7%) | IS: 716 (0.3%) |
| On blood-pressure lowering<br>medication, N (%) | 80,880 (20.5%) | 43,714 (24.2%) | 37,166 (17.3%) |
| On lipid-lowering medication, N<br>(%) | 66,739 (16.9%) | 40,164 (22.2%) | 26,575 (12.4%) |
| Follow-up time, years [mean (sd)] | 6.3 (1.9) | 6.2 (2.1) | 6.4 (1.8) |

We used GWAS summary statistics that did not include the UKB for five stroke outcomes and 14 stroke-related phenotypes (**Supplementary Table 1**) to generate 19 GRSs associated with ischaemic stroke (**Figure 1**). As expected, the 19 individual GRSs were correlated with each other in several distinct clusters: (i) any stroke (AS), ischaemic stroke (IS), cardioembolic stroke (CES), large artery stroke (LAS), and small vessel stroke (SVS); (ii) the three CAD scores (1KGCAD, 46K, and FDR202); (iii) total cholesterol (TC), triglycerides (TG), low density lipoprotein cholesterol (LDL), and high density lipoprotein cholesterol (HDL); (iv) systolic blood pressure (SBP) and diastolic blood pressure (DBP); and (v) body mass index (BMI) and type 2 diabetes (T2D) **(Figure 2**). From the 19 distinct GRSs, we constructed the metaGRS using elastic-net logistic regression with 10-fold cross-validation on the derivation set (**Figure 1**; metaGRS; model weights are shown in **Supplementary Figure 1**), and subsequently converted the model to a set of 3.2 million SNP weights, which have been made freely available (https://dx.doi.org/10.6084/m9.figshare.8202233).

**Figure 2:**
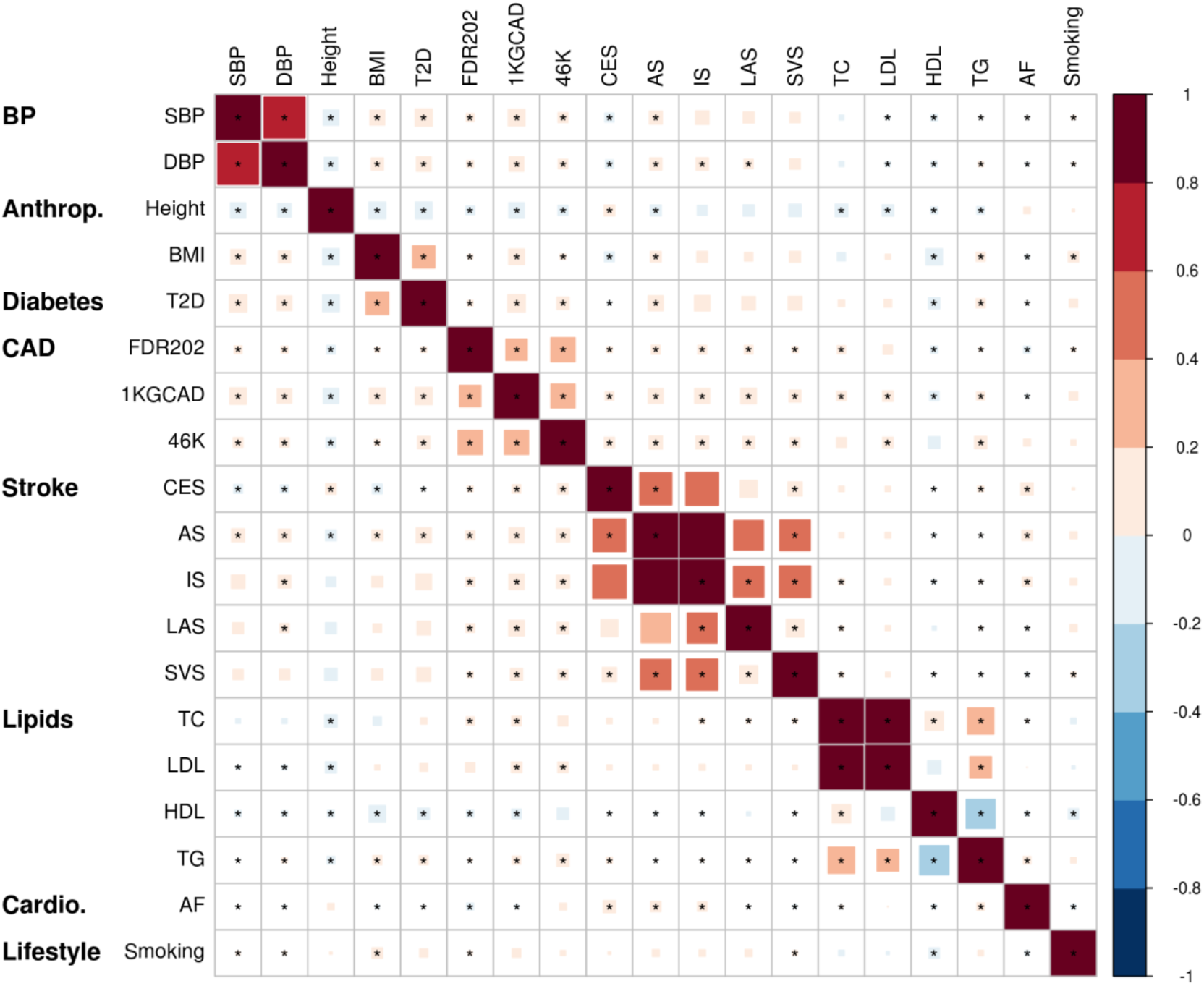
Individual GRSs for stroke-related phenotypes and stroke outcomes correlate in several distinct clusters. Shown is the correlation plot of individual GRSs in a random sample of 20,000 UK Biobank individuals. Estimates are from linear regression of each pair of standardised GRSs, adjusting for genotyping chip (UKB/BiLEVE) and 10 PCs. Stars indicate Benjamini-Hochberg false discovery rate <0.05 (adjusting for 171 tests). GRSs were ordered via hierarchical clustering of the absolute correlation. *Anthrop*, anthropometric*; cardio:* cardiovascular (other than CAD)*; SBP*: systolic blood pressure; *DBP*: diastolic blood pressure; *Height*: measured height; *BMI*: body mass index; *T2D*: type 2 diabetes; *1KGCAD*: coronary artery disease from 1000 Genomes; *46K*: coronary artery disease from Metabochip; FDR202: coronary artery disease from 1000 Genomes (top SNPs); *CES*: cardioembolic stroke; *AS*: any stroke; *IS*: ischaemic stroke; *LAS*: large artery stroke; *SVS*: small vessel stroke; *TC*: total cholesterol; *LDL*: low density lipoprotein cholesterol; *HDL*: high density lipoprotein cholesterol; *TG*: triglycerides; *AF*: atrial fibrillation; *Smoking*: cigarettes per day.

### The metaGRS improves risk prediction of ischaemic stroke compared to other genetic scores

Using the independent UKB validation set, we next quantified the risk prediction performance of the metaGRS, and evaluated its association with IS via survival analysis. The metaGRS was associated with IS with a HR of 1.26 (95% CI 1.22-1.31) per standard deviation of metaGRS, which was stronger than any individual GRS comprising the metaGRS and was twice the effect size of the previously published 90-SNP IS GRS^24^ (HR=1.13 [95% CI 1.10-1.17]; **Supplementary Figure 2a**). The metaGRS also increased the C-index by 0.029 over the 90-SNP GRS (**Supplementary Figure 2b**). We also assessed the performance of the IS metaGRS for predicting the any stroke (AS) outcome. We found the associations were consistently weaker for AS than for IS, however, as with IS, the metaGRS was a stronger predictor of AS than the 90-SNP GRS score (**Supplementary Figure 2**).

In a Kaplan-Meier analysis of IS, the top and bottom 10% of the metaGRS showed substantial differences in cumulative incidence of IS (**Supplementary Figure 3**; log-rank test between the top decile and the 45-55% decile: P=3×10^−6^); these results were consistent with a Cox proportional hazards model of the metaGRS assessing the HRs for the top 10% decile vs the middle 45-55% decile (**Supplementary Figure 4**). The top 0.25% of the population were at a three-fold increased risk of IS versus the middle decile (45-55%), with HR=3.0 (95% CI 1.96-4.59) (**Figure 3**).

**Figure 3:**
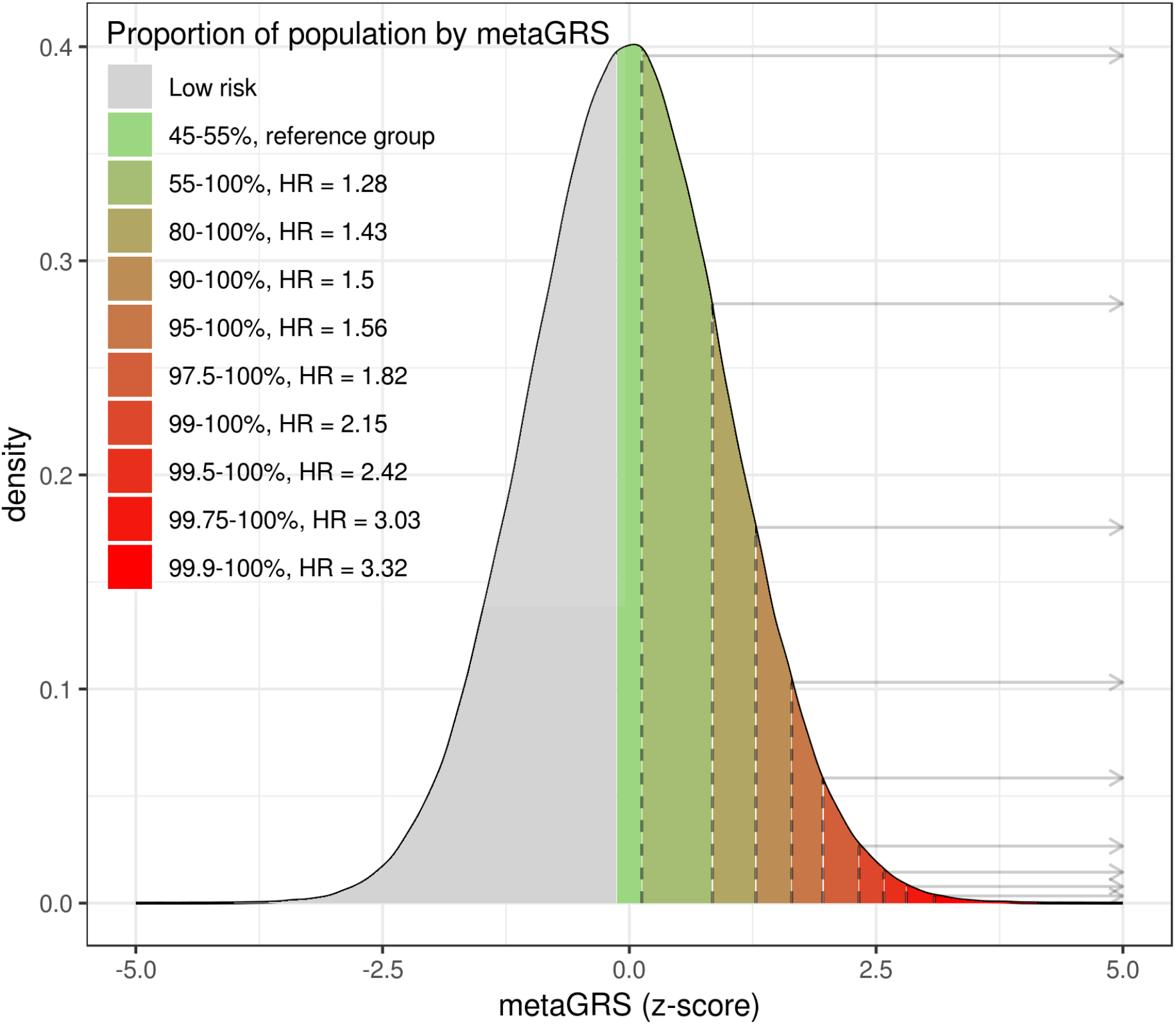
The metaGRS identifies individuals at increased risk of ischaemic stroke. Shown is the distribution of the metaGRS for ischaemic stroke in the UK Biobank validation set, and corresponding hazard ratios. Hazard ratios are for the top metaGRS bins (stratified by percentiles) versus the middle metaGRS bin (45-55%).

There was no evidence for a statistical interaction of the metaGRS with sex on IS hazard (P=0.614), indicating that the substantial differences in cumulative incidence between the sexes were driven by differences in baseline hazards rather than by any sex-specific effects of the metaGRS itself.

### The ischaemic stroke metaGRS has comparable or higher predictive power than established risk factors

We next compared the performance of the metaGRSs with established risk factors^31^ for predicting IS. We examined seven established risk factors at first UKB assessment: LDL cholesterol, SBP, family history of stroke, BMI, diabetes diagnosed by a doctor, current smoking, and hypertension (an expanded definition based on SBP/DBP measurements, BP medication usage, self-reporting, and hospital records; **Methods**).

As expected, established risk factors were positively associated with incident IS, with hypertension being the strongest risk factor (**Supplementary Figure 5**). Notably, the HR of the metaGRS (incident IS HR=1.25 per s.d.) was similar to that of SBP (incident IS HR=1.28 per s.d., where the s.d. of SBP was 21.7mmHg) and current smoking (incident IS HR=1.25, s.d.=0.3) (**Supplementary Figure 5)**.

Comparison of the C-index for time to incident IS revealed that blood pressure phenotypes, hypertension and SBP (C=0.590 [95% CI 0.577-0.603]; C=0.584 [95% CI 0.570-0.598], respectively), had the largest C-indices followed by the metaGRS (C=0.580 [95% CI 0.566-0.593]) and the other established risk factors (**Figure 4**). Notably, the metaGRS had a greater C-index than family history of stroke (C=0.558, 95% CI 0.544-0.572; **Figure 4**). The metaGRS and hypertension contained similar additional information on top of the other risk factors; adding either the metaGRS or hypertension to the six other risk factors yielded similar predictive power, C=0.629 (95% CI 0.615-0.643) and C=0.628 (95% 0.614-0.641), respectively. Finally, adding both the metaGRS and hypertension to the six risk factors yielded the model with the highest C-index, C=0.637 (95% CI 0.623-0.650) (**Figure 4**).

**Figure 4:**
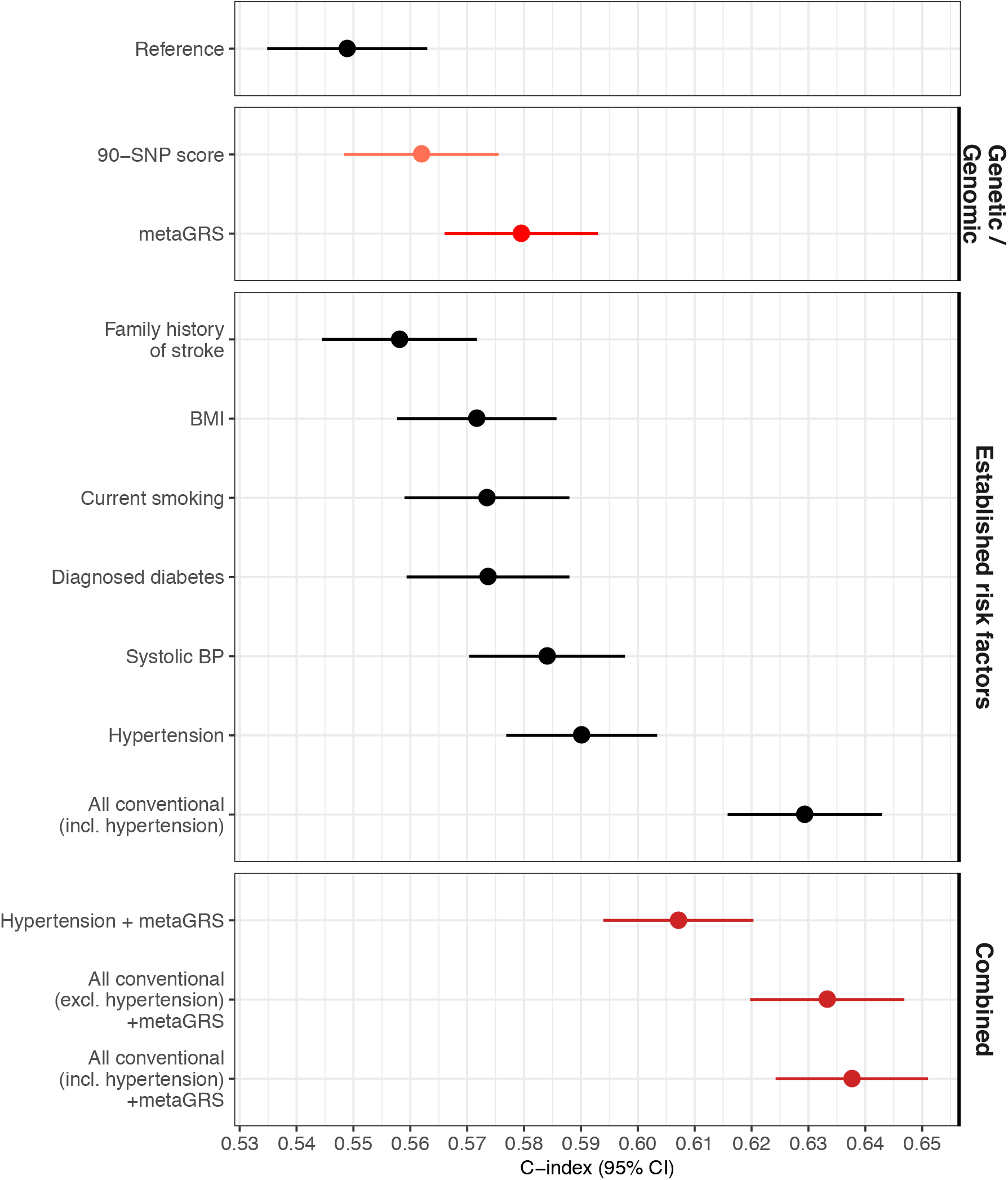
The metaGRS for ischaemic stroke has comparable or higher predictive power than established risk factors. Shown are the C-indices for incident stroke in the UKB validation set comparing the metaGRS with established risk factors. The reference model included the genotyping chip and 10 genetic PCs.

### The metaGRS contributes to ischaemic stroke risk independent of established risk factors

Given that the metaGRS is composed of GRSs for stroke and stroke risk factors, we conducted several complementary analyses to assess the association of the metaGRS with these risk factors, and whether the metaGRS was associated with IS risk independently of these risk factors. As expected, the IS metaGRS was positively and significantly associated with all seven risk factors (**Supplementary Table 2**). Adjusting for these risk factors as well as BP-lowering and/or lipid-lowering medication status only modestly attenuated the association of the metaGRS with incident IS (**Supplementary Figure 6**), indicating that the information contained in the metaGRS was only partially explained by these factors. On the other hand, adjusting for the metaGRS modestly but consistently attenuated the association of each risk factor itself with IS risk (**Supplementary Figure 5**). There was no evidence for statistical interaction of the metaGRS effects on IS with medication status at assessment (logistic regression, P=0.23 and P=0.82 for interaction of the metaGRS with BP medication and cholesterol-lowering medication, respectively).

### Predicting ischaemic stroke risk with established risk factors and the metaGRS

The clinical utility of a GRS depends on its performance in combination with established risk factors and risk models. To examine this, we conducted analyses integrating information on risk factor levels based on (i) recent ACC / AHA / AAPA / ABC / ACPM / AGS / APhA / ASH / ASPC / NMA / PCNA guidelines^32^ (SBP<120 mmHg); (ii) AHA / ASA guidelines for primary prevention of stroke^31^ (BMI<25 kg/m^2^); (iii) smoking status and diabetes status. We used Cox models of these established risk factors and the metaGRS together with the estimated baseline cumulative hazards to predict cumulative incidence of IS for individuals with a high metaGRS (top 1%), average metaGRS (50%), and low metaGRS (bottom 1%) along with two levels of risk factors: (i) meeting guideline targets for the above risk factors^32^ and (ii) the following combination of risk factors representative of an individual at typical stroke risk: SBP=140 mmHg, BMI=30 kg/m^2^, current smoking, and no diagnosed diabetes.

The predicted risk of IS for individuals with a high metaGRS (top 1%) and high levels of risk factors was maximal by age 75, reaching a cumulative incidence of 8.5% (95% CI 5.2-11.6%) for males and 5.1% (95% CI 3.1-7.1%) for females (**Figure 5a**). Effective reduction in the levels of the modifiable risk factors (SBP, BMI, and smoking) to match guideline targets was predicted to result in a substantial reduction in risk, down to 2.8% (95% CI 1.7-3.9%) for males and 1.7% (95% CI 1.0-2.4%) for females by age 75, thus substantially compensating for the high genomic risk.

**Figure 5:**
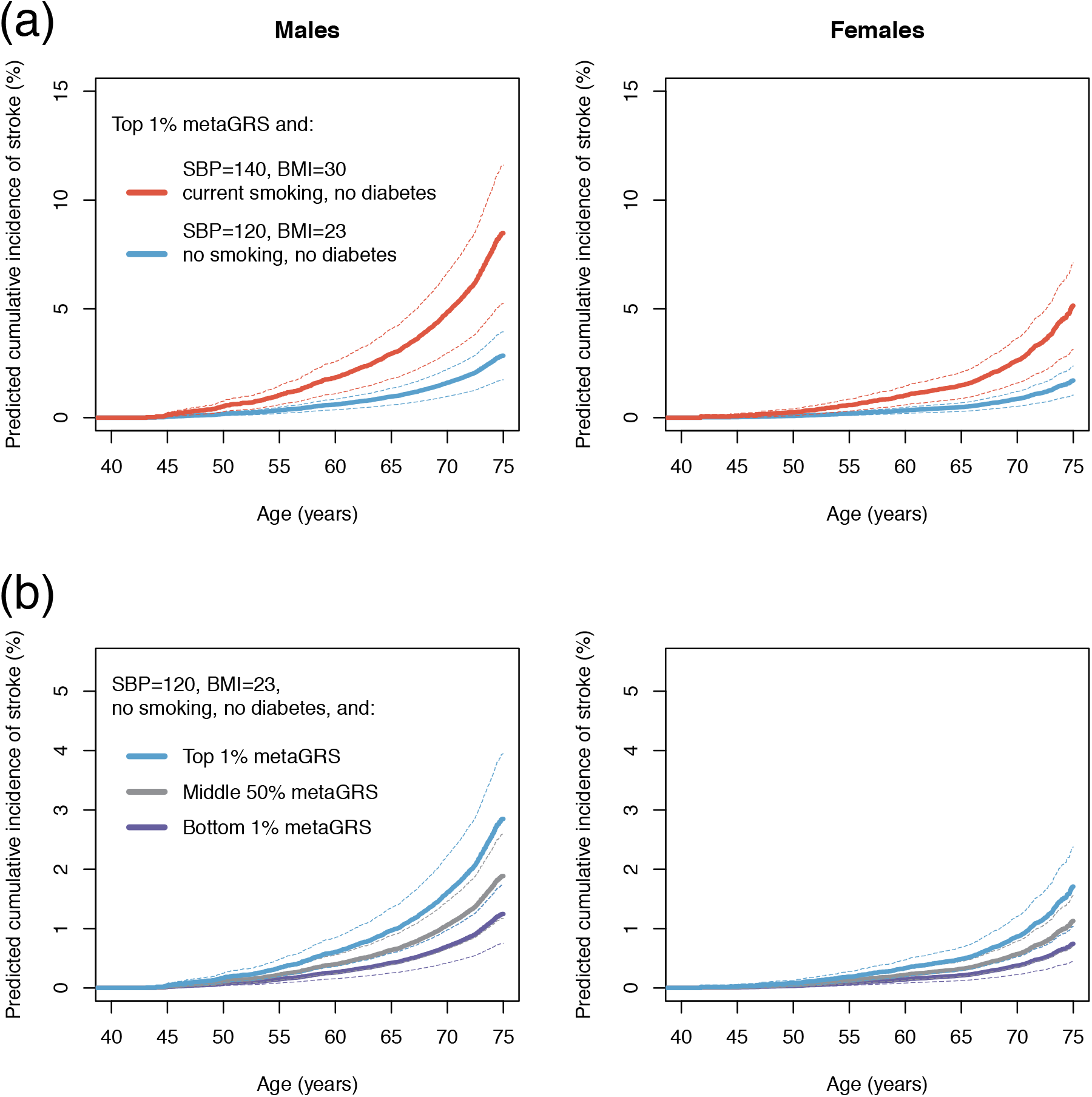
Predicted cumulative incidence of ischaemic stroke. Shown is the predicted cumulative incidence of IS in subjects with either (a) high levels of the metaGRS along with different risk factor levels (within or outside the guidelines); or (b) risk factors within accepted guidelines along with different levels of the metaGRS.

Conversely, for individuals matching the guidelines for established risk factors (**Figure 5b**), there were notable differences in IS incidence for individuals in the top (1%) compared to the bottom (1%) of the metaGRS; with 2.8% (95% CI 1.7-3.9%) versus 1.2% (95% CI 0.7-1.7%) in males and 1.7% (95% CI 1.0-2.4%) versus 0.7% (95% CI 0.4-1.0%) in females, respectively, by age 75. These results further indicate that the metaGRS captures residual risk of stroke not quantified by existing risk factors.

## Discussion

In this study, we developed a genomic risk score for ischaemic stroke based on GWAS summary statistics for 19 stroke and stroke-related traits. We quantify the predictive power of the IS metaGRS by comparing it to previously published genetic scores and measures of established non-genetic risk factors, and demonstrate its added value in combination with established risk factors and in the context of current guidelines for primary stroke prevention. While genomic risk scores for stroke are not yet at the level necessary for clinical translation, our analyses constitute several significant advances.

First, we showed that the IS metaGRS had stronger association with IS than previously published genetic scores, doubling the effect size of the most recent genetic score. To put its performance in context, we estimated the IS metaGRS identified the 1 in 400 individuals who were at 3-fold increased risk of IS, a level of risk and frequency similar to common monogenic cardiovascular diseases, such as familial hypercholesterolemia (FH), a risk factor for myocardial infarction^33^. Monogenic forms of stroke, such as CADASIL, are relatively rare^34^, thus the IS metaGRS may represent a potential new avenue to more common polygenic risk stratification, in combination with established risk factors.

Second, the IS metaGRS had comparable predictive power to systolic blood pressure, and higher predictive power than other established risk factors measured, apart from hypertension, and captured residual risk not quantified by the established risk factors. In anticipation of a potential role in early screening, we estimate the risk reduction through modifiable stroke risk factors across different metaGRS backgrounds, and further show that current guidelines for stroke risk factors may be insufficiently stringent for individuals at high metaGRS.

Third, we explicitly modelled how changes in modifiable risk factors, such as systolic blood pressure and body mass index, can compensate for high genomic risk. Previous research has demonstrated that intervening on modifiable risk factors can compensate for increased genetic risk of disease^21,35^. However, these analyses relied on simply counting the number of elevated risk factors, which does not account for the differences in effect size between various risk factors. Importantly, our approach was flexible in that various combinations of risk factor reductions can lead to the same outcome in terms of risk.

Our approach shows, for different genomic risk backgrounds, how modifiable risk factors could, in principle, be tailored to an individual’s ability to reduce an established risk factor(s) while maintaining an overall acceptable level of absolute risk. Similarly, this approach could potentially be used to guide early prevention of stroke: identifying individuals at increased risk early in life, who would then be targeted for more intensive lifestyle modifications, similar to the roles that have been proposed for genetics in cancer risk stratification^36^. Unlike most established risk factors which may vary over time and are typically not informative at an early age, the metaGRS remains stable and can be derived from birth. Later in life, when measurements of established risk factors are available, these can be further combined with the metaGRS to give the most accurate prediction of a person’s risk of incident stroke. Further research is required to determine what levels of risk factor reductions will be achievable and cost effective in practice.

Lastly, even for individuals within risk factor levels recommended by current guidelines (SBP<120, BMI<25, not currently smoking, no diagnosed diabetes), our models predict substantial differences in risk between different metaGRS levels. These results suggest that for individuals with high metaGRS, achieving currently recommended risk factor levels may not be sufficient and that it is time to contemplate whether future guidelines on primary and secondary stroke prevention should integrate genetic information when defining treatment goals for high risk individuals.

Our study has several limitations. The GWAS for stroke are still themselves limited by phenotypic heterogeneity and are less powered compared to other common diseases, such as CAD^21^. As stroke GWAS progress, genomic risk scores will become more powerful^37,38^. We did not observe substantial advantage from incorporating GRSs based on GWAS summary statistics for specific IS subtypes (LAS, CES, SVS) over that of IS as a whole. However, there may be benefit from developing subtype-specific scores that take advantage of the unique genetic architecture of each subtype^39^. The number of older individuals (>75 years) in UKB is limited at this stage, reducing our ability to model stroke risk in the age strata where the majority of events occur. Furthermore, the duration of follow-up in UKB is relatively limited and, because of the limited number of assessments, we could not model the cumulative effect of BP and smoking over time; however, we accounted for potential regression dilution bias in SBP measurements via the use of diagnosed hypertension, which showed stronger associations with stroke. Family history of stroke in UKB may be less comprehensive than in stroke-specific studies, limiting its predictive power, and overall the UKB study population is healthier than the general UK population^40^, which could have led to underestimation of some of the effects of risk factors. Our modelling assumes that risk factors, such as SBP and BMI, can be varied independently of each other. In practice, common lifestyle interventions such as exercise and diet will likely affect several risk factors at a time. Finally, this study focused on British white ancestry, and further studies are required to validate these scores in other populations^41^.

Taken together, despite challenges in phenotypic heterogeneity and corresponding GWAS power, our study presents the most powerful ischaemic stroke genomic risk score to date and assesses its potential for risk stratification in the context of established risk factors and clinical guidelines. It lays the groundwork for larger GWAS of stroke and its multiple sub-types as well as analyses which leverage the totality of information available for stroke genomic risk prediction.

## Methods

### Study participants

The UK Biobank (UKB) study^29,30^ included individuals from the general UK population, aged between 40-69 years at recruitment. Recruitment included a standardised socio-demographic questionnaire, as well as medical history, family history, and other lifestyle factors. Several physical measurements (e.g., height, weight, waist-hip ratio, systolic and diastolic BP) were taken at assessment.

Individual records were linked to the Hospital Episode Statistics (HES) records and the national death and cancer registries. The age of event was age at the primary stroke event (the diagnostic algorithm for stroke in UKB can be found at http://biobank.ndph.ox.ac.uk/showcase/docs/alg_outcome_stroke.pdf; last accessed 11/04/2019).

We defined stroke risk factors at the first assessment, including: diabetes diagnosed by a doctor (field #2443), body mass index (BMI; field #21001), current smoking (field #20116), hypertension, family history of stroke, and high cholesterol. For hypertension we used an expanded definition including self-reported high blood pressure (either on blood pressure medication, data fields #6177, #6153; or systolic blood pressure >140 mmHg, fields #4080, #93; or diastolic blood pressure >90 mmHg, data fields #4079, #94) as well as hospital records; for registry cases, we use HESIN (hospital admission) and death registry data including both primary and secondary diagnoses / causes of death (HESIN: ICD9 401-405, ICD10 I10-I15; death: ICD10 I10-I15, data fields #40001, #40002). For family history of stroke, we considered history in any first degree relative (father, mother, sibling; fields #20107, 20110, and 20111, respectively).

We excluded individuals with withdrawn consent, self-reported stroke at age <20 years due the potential unreliability of these records, and those not of British white ancestry (identified via the UKB field ‘in.white.British.ancestry.subset’^29^), leaving a total of n=407,388 individuals. We censored the age of stroke at 75y.

For individuals on BP-lowering medication, we adjusted systolic blood pressure by adding +15mmHg as per Evangelou et al^16,42^. We used LDL-cholesterol from the UKB biomarker panel, measured at first UKB assessment. For individuals on lipid-lowering medication at the time of assessment (n=66,737), we adjusted the measured LDL-cholesterol level by +1.5 mmol/L.

### Genotyping quality control

The UKB v2 genotypes were genotyped on the UKB Axiom array, and imputed to the Haplotype Reference Consortium (HRC) by the UKB^29^; SNPs on the UK10K/1000Genomes panel were excluded from the current analysis. Imputed genotypes were converted to PLINK hard calls. For the initial GRS analysis, we considered genotyped or HRC-imputed SNPs with imputation INFO >0.01 and global MAF >0.001 (14.5M autosomal SNPs). A further QC step was performed on the final metaGRS (see below).

### Generation of the metaGRS

We randomly sampled n=11,995 individuals from the UKB dataset, oversampling individuals with any stroke (AS) events, leading to 2065 individuals with AS (of which 889 were also IS events) and 9935 non-AS referents. This subset was used for developing GRSs, and was excluded from all further analysis. Five individuals were later removed due to withdrawn consent.

Using the UKB derivation set, we generated 19 GRSs for phenotypes associated with stroke (**Supplementary Table 1**). To minimise the risk of over-fitting due to inclusion of the same individuals in the derivation and validation datasets we selected GWAS that did not include the UK Biobank in their meta-analysis.

The three CAD GRSs (46K, 1KGCAD, FDR202) were generated previously using an n=3000 derivation subset of the UKB (included in the larger n=12,000 subset employed here)^21^. The AF GRS was defined previously^43^. For the remaining GRSs, we used published summary statistics to generate a range of scores based on different *r*^2^ thresholds with PLINK ^44^ LD thinning (--indep-pairwise), and selected one optimal model (in terms of the largest magnitude hazard ratio), resulting in one representative GRS for each set of summary statistics.

Each GRS was standardised (zero mean, unit standard deviation) over the entire dataset. Next, we employed elastic-net logistic regression^45^ using the R package ‘glmnet’^46^ to model the associations between the 19 GRSs and stroke, adjusting for sex, genotyping chip (UKB vs BiLEVE), and 10 genetic PCs. A range of models with different penalties were evaluated using 10-fold cross-validation. The best model, in terms of highest cross-validated AUC (area under receiving-operator characteristic curve), was selected as the final model and held fixed for validation in the rest of the UKB data. The final adjusted coefficients for each GRS (odds ratios) in the penalised logistic regression are shown in **Supplementary Figure 1**, in comparison with the univariate estimates (based on one GRS at a time).

The final per-GRS log odds *γ*_1_, …, *γ*_19_ were converted to an equivalent per-SNP score via a weighted sum

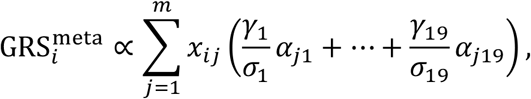

Where *m* is the total number of SNPs, *σ*_1_, …, *σ*_19_ are the empirical standard deviations of each of the 19 GRSs in the derivation data, *α*_*j*1_, …, *α*_*j*19_ are the SNP effect sizes (from the GWAS summary statistics) for the *j*th SNP in each of the GRSs, respectively, and *x*_*ij*_ is the genotype {0, 1, 2} for the *i*th individual’s *j*th SNP. A SNP’s effect size *α*_*jk*_ was considered to be zero for the *k*th score if the SNP was not included in that score. This resulted in 3.6 million SNPs for inclusion in the metaGRS.

We conducted a sensitivity analysis to evaluate whether stricter quality control filtering would impact the performance of the metaGRS; removing SNPs with imputation INFO<0.4 and MAF<0.01 did not substantially affect the association of the metaGRS with stroke, hence, we selected the metaGRS with stricter QC as the final score, bringing the total number of SNPs to 3.2 million.

### Evaluation of the metaGRS

The metaGRS developed using the derivation set was held fixed and evaluated in the UKB validation subset (n=395,393) using a Cox proportional hazard model. We conducted complete case analysis due to the low proportion of participants with any missing values for the seven risk factor variables of interest (5.1% of participants).

Age was used as the time scale in the Cox proportional hazard regression. The regression was stratified by sex and weighted by the inverse probability of selection into the validation set, together with robust standard errors (R package ‘survival’^47^). All analyses were adjusted for chip (UKB vs BiLEVE) and 10 PCs of the genotypes (as provided by UKB^29^). For analyses of incident stroke, age at UKB assessment was taken as time of entry into the study. Cox models of the metaGRS did not show deviations from proportional hazard assumptions, based on the global test for scaled Schoenfeld residuals (P=0.32).

The predicted cumulative risk curves (as a function of time *t*) were calculated using ‘survfit.coxph’ within each stratum of sex as

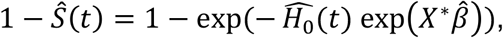

where 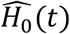 is the estimated baseline cumulative hazard, *X*^*^ is the matrix of the predictor variables set to the values of interest and 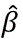 is the vector of the estimated log hazard ratios.

We performed a sensitivity analysis testing whether the association of the metaGRS with ischaemic stroke was affected by familial relatedness in the validation set. Relatedness analysis was done using KING^48^ v2.1.4, based on ~784,000 autosomal SNPs measured on the Axiom chip, identifying n=336,643 participants in the UKB validation set with kinship more distant than that of 2^nd^ degree. There was a negligible difference in the association between the metaGRS and stroke in the full UKB validation set and within this distantly-related subset of individuals.

Calibration of the metaGRS risk score was evaluated by fitting logistic regression models of the metaGRSs (adjusting for sex, chip, and 10 genetic PCs) in the derivation set, predicting the absolute risk of event in the test set (allowing for the 9.38-fold lower observed baseline rate of events between the testing set compared with the derivation set), and evaluating the proportion of test-set individuals with stroke events within each decile of the predicted risks (**Supplementary Figure 7**). Pointwise confidence intervals were obtained via the binomial test for proportions.

We estimated the heritability of ischaemic stroke explained by the metaGRS, on the liability scale, using the *R*^2^ and partial *R*^2^ obtained from linear regression of the stroke outcomes on metaGRS (partial *R*^2^ was from linear regression adjusted for sex, age of assessment, genotyping chip, and 10 PCs). The estimates were converted to the liability scale^49^, assuming that the ischaemic stroke prevalence in UKB represents that of the general population (*K*=0.008). Due to a lack of robust estimates of the heritability of stroke, we examined a range of plausible *h*^2^ values from 0.1 to 0.4, yielding estimates of explained heritability ranging from 7.7% to 1.8%, respectively (**Supplementary Figure 8**).

We performed sensitivity analysis to assess the effect of potential geographical stratification within the UKB^50^ on the metaGRS. We compared the original metaGRS with residuals of the metaGRS regressed on (i) the first 10 PCs, (ii) first 10 PCs and natural cubic splines of the geographical north-south coordinate and east-west coordinates (3 degrees of freedom each), (iii) the first 30 PCs and splines of the coordinates, (iv) first 10 PCs and a thin-plate regression spline (TPRS) representing smooth interactions between the two coordinates^51^, and finally (v) also adding the UKB assessment centre (**Supplementary Figure 9a**). For the unadjusted score, we observed some variation across the north and east coordinates (up to 0.4 standard deviations), however, adjusting for PCs and the coordinates attenuated this variation substantially, with the TPRS method eliminating it completely. Despite the attenuation in geographical stratification, we observed negligible change in the association of the residuals of the scores with IS events (**Supplementary Figure 9b**), indicating that any geographical stratification in UKB was not driving the metaGRS’s association with stroke.

## Acknowledgments

This study was supported in part by the Victorian Government’s OIS Program. MI was supported by an NHMRC and Australian Heart Foundation Career Development Fellowship (no. 1061435). GA was supported by an NHMRC Early Career Fellowship (no. 1090462). MD acknowledges funding from the European Union’s Horizon 2020 research and innovation programme (grant agreements No 666881, SVDs@target and No 667375, CoSTREAM); and the DFG as part of the Munich Cluster for Systems Neurology (EXC 1010 SyNergy) and the CRC 1123 (B3); The MEGASTROKE project received funding from sources specified at http://www.megastroke.org/acknowledgments.html. Data on coronary artery disease / myocardial infarction have been contributed by CARDIoGRAMplusC4D investigators and have been downloaded from www.cardiogramplusc4d.org. The MRC/BHF Cardiovascular Epidemiology Unit is supported by the UK Medical Research Council [MR/L003120/1], British Heart Foundation [RG/13/13/30194], and UK National Institute for Health Research Cambridge Biomedical Research Centre. Dr. Butterworth has received grant support from Merck, Novartis, Pfizer, Biogen, Bioverativ, and AstraZeneca; and serves as a consultant to Novartis.

UK Biobank analyses were conducted under project 26865.

