## Supplementary Figures and Tables for "Genomic risk score offers predictive performance comparable to clinical risk factors for ischaemic stroke"

**Figure S1: Associations of individual GRSs with ischaemic stroke in the UKB derivation set;** Shown are the odds ratios per standard deviation of each GRS estimated in either (i) logistic regression of each GRS, adjusting for chip (UKB/BiLEVE), 10 PCs, and sex; or in (ii) elastic-net logistic regression. The elastic-net estimates are from the best model selected via cross-validation, adjusting for all other GRSs, chip, sex, and 10 genetic PCs. For elastic-net, confidence intervals are not available; 'inactive' indicates that the elastic-net estimated odds ratio was negligible (between 0.999 and 1.001). Scores are ordered by their univariate associations with ischaemic stroke.

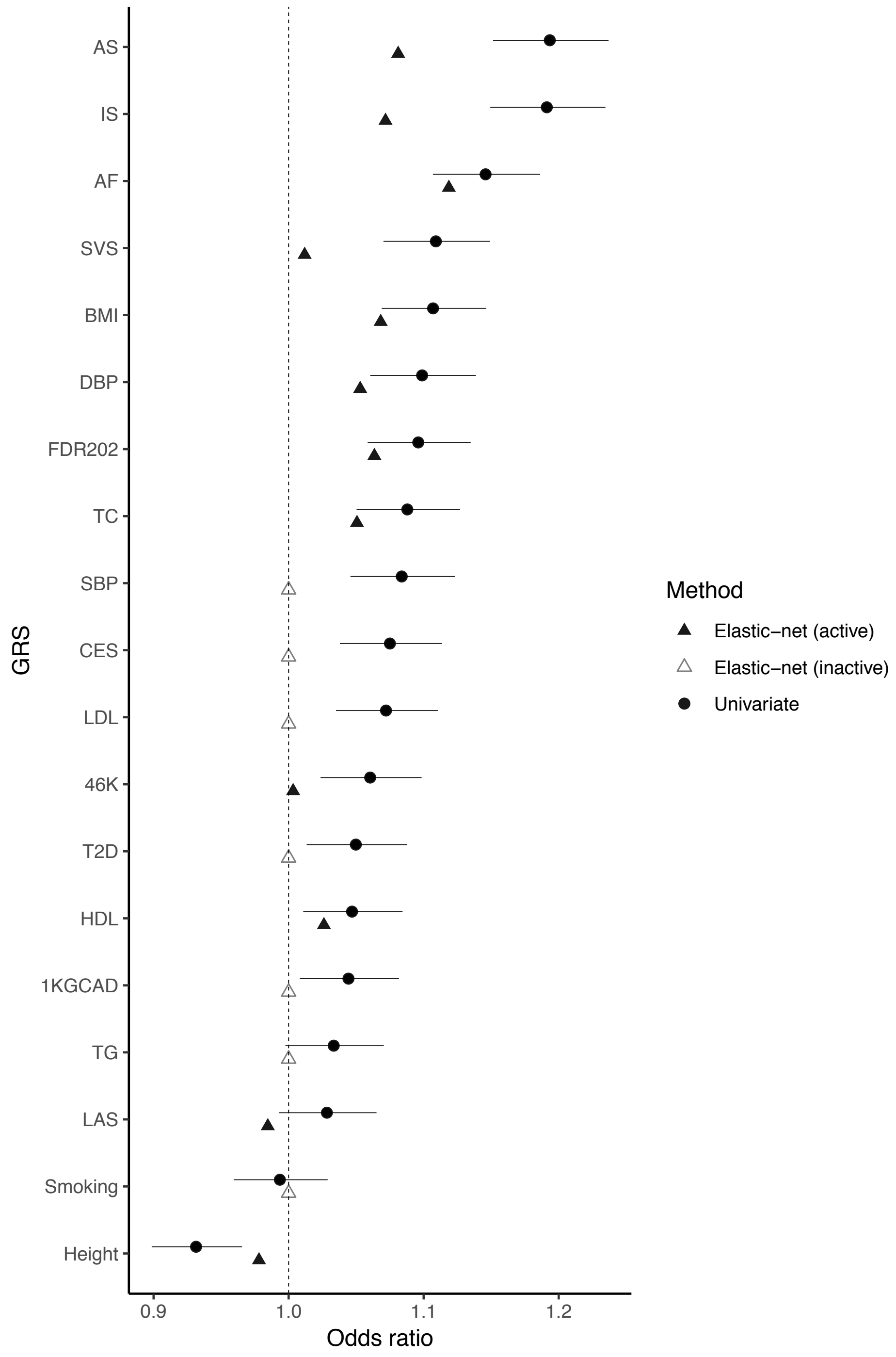

**Figure S2: Comparison of the metaGRS with a published 90-SNP risk score for stroke[20] and the ischaemic stroke GRS derived earlier, predicting both prevalent and incident any stroke (AS) and ischaemic stroke (IS), in the UK Biobank validation set.** Scores were standardised to zero-mean, unit standard deviation. Results are from Cox regression on the UKB validation set (n=395,393), sex-stratified, adjusted for chip and 10 PCs.

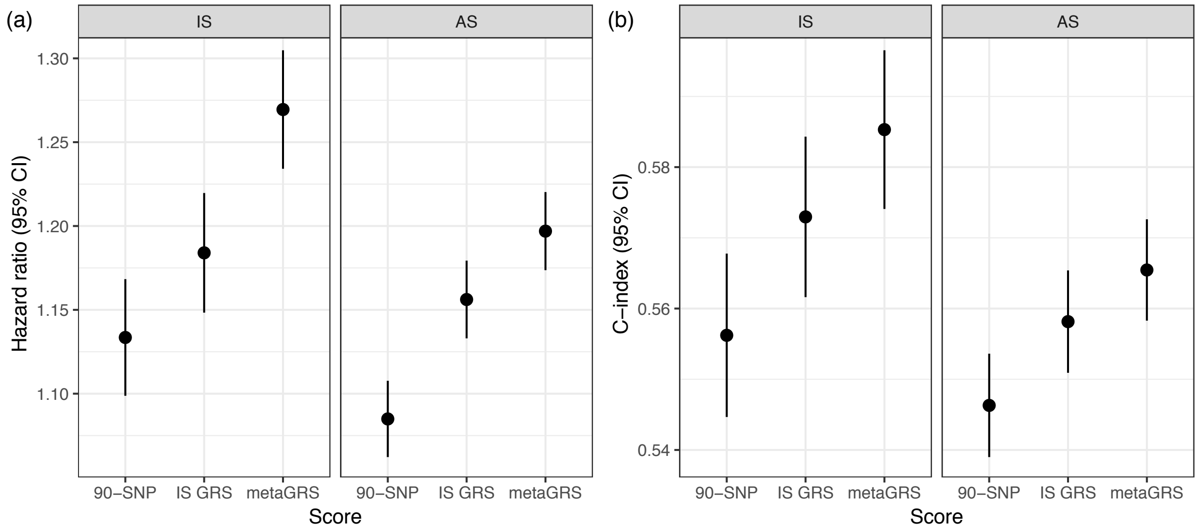

**Figure S3: Cumulative incidence of ischaemic stroke stratified by the metaGRS (prevalent and incident stroke) in the UK Biobank validation set.** From age-as-time-scale Kaplan-Meier analysis.

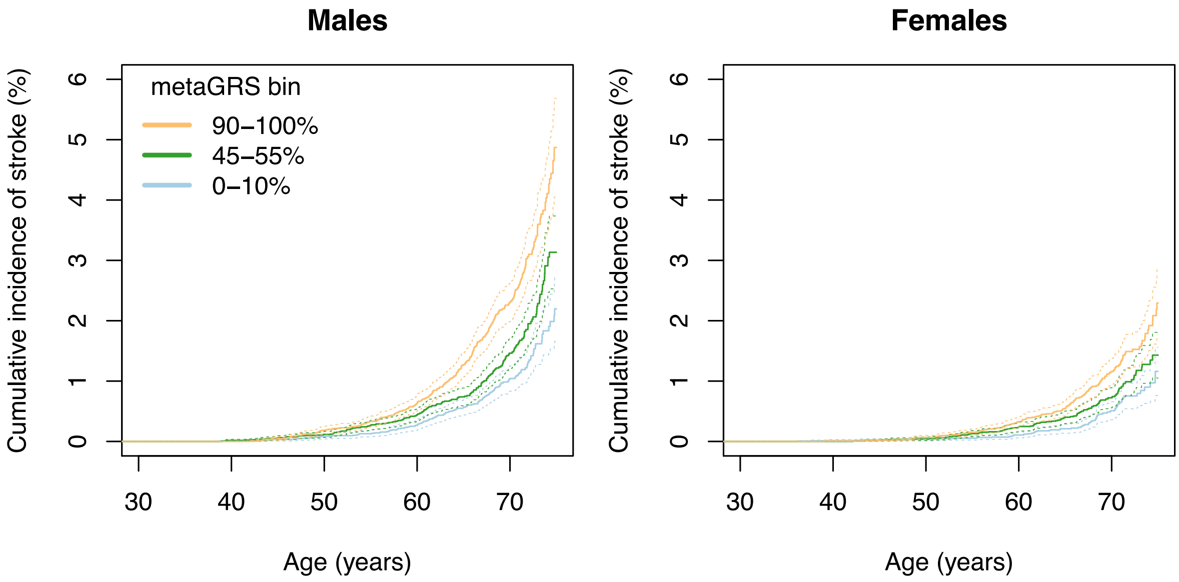

**Figure S4: Hazard ratios for different quantiles of the metaGRS (relative to the 45–55 centiles of the metaGRS distribution) for all recorded ischaemic stroke (prevalent and incident) in the UK Biobank validation set.** Based on age-as-time-scale Cox regression stratified by sex and adjusted for chip and 10 genetic PCs.

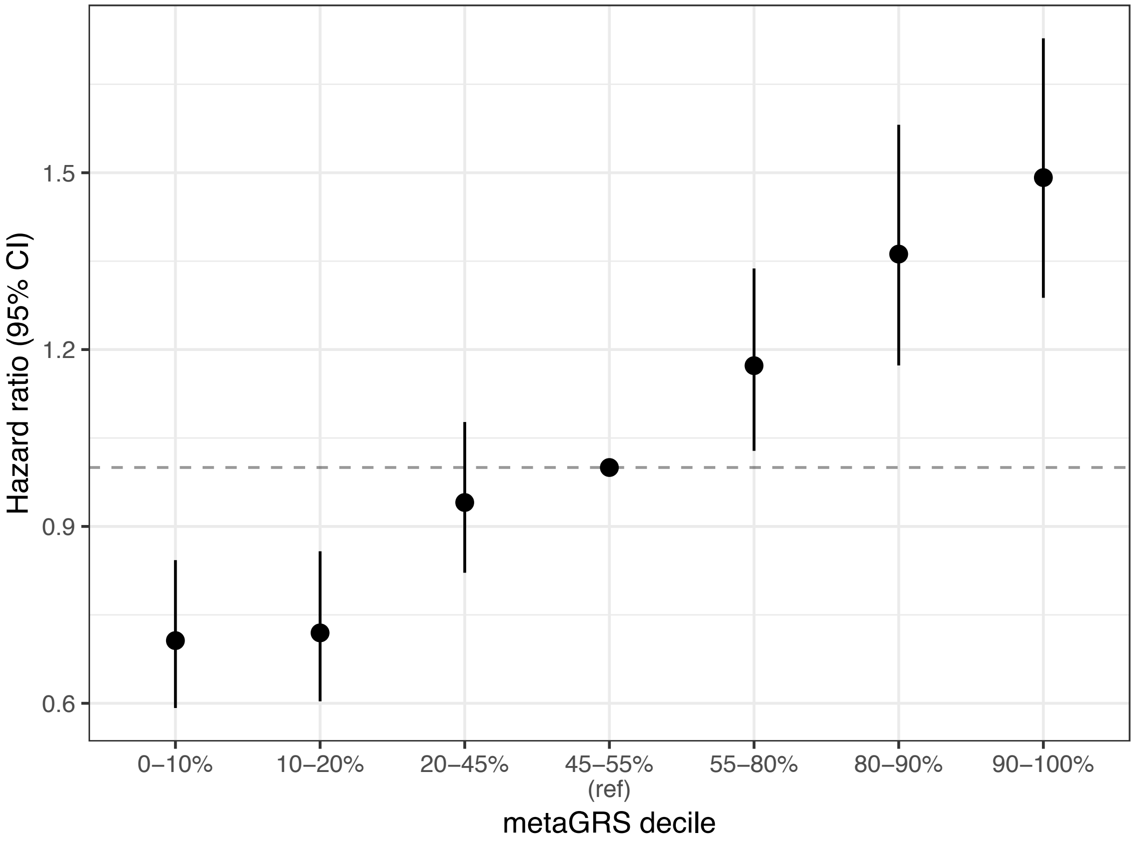

**Figure S5: Association of established risk factors with incident ischaemic stroke, adjusted or not adjusted for the metaGRS.** Hazard ratios (95% CI) are from Cox regression models of each risk factor (with or without the metaGRS), stratified by sex and adjusting for chip and 10 PCs. Results are presented per standard deviation of each factor.

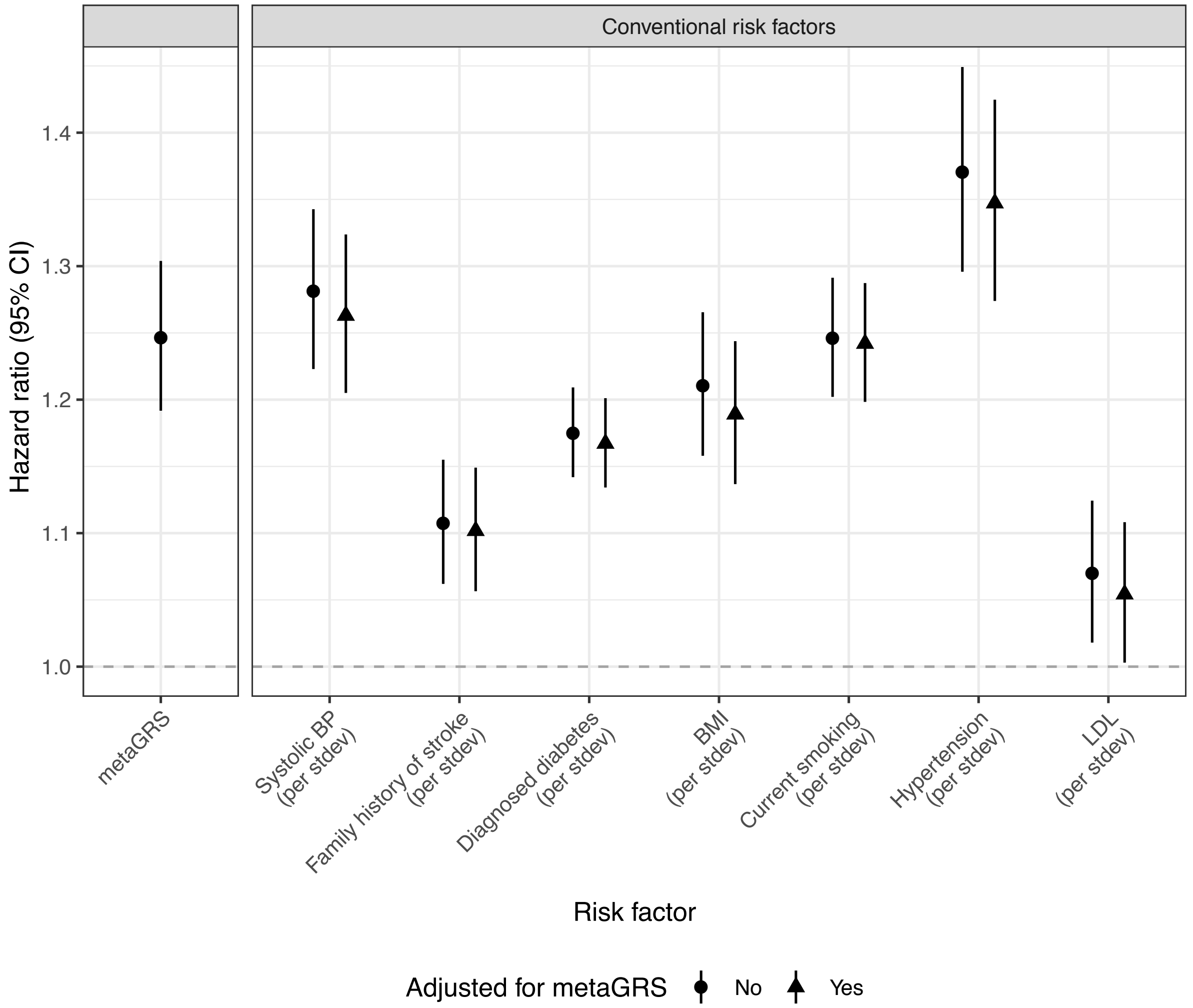

**Figure S6: Hazard ratios of incident ischaemic stroke for the metaGRS in the UK Biobank validation set.**

(1) ‘*metaGRS*’: sex-stratified, adjusted for chip and 10 PCs;
(2) ‘*Chol medication + metaGRS*’: adjusting for cholesterol-lowering medication status;
(3) ‘*BP medication +. metaGRS*’: adjusting for BP medication status;
(4) ’*BP+chol medication+metaGRS*’: adjusting for both cholesterol and BP lowering medications;
(5) ‘*All conventional+metaGRS*’: adjusting for established risk factors (systolic BP, diastolic BP, family history of stroke, body mass index, current smoking, diagnosed high cholesterol, hypertension);
(6) ‘*All conventional+medication+metaGRS*’: adjusting for all conventional risk factors as well as cholesterol-lowering and BP medication.

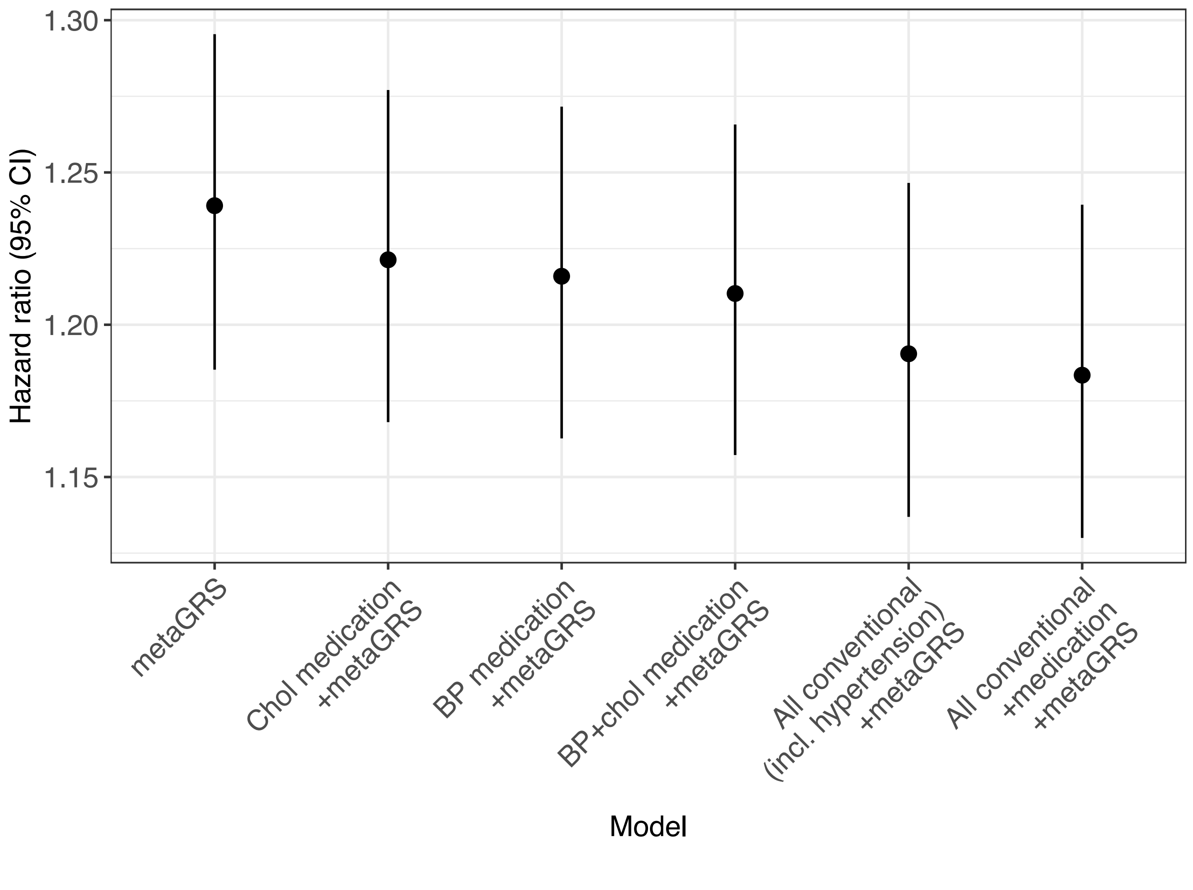

**Figure S7: Calibration of logistic regression models of the metaGRS in the UK Biobank validation set (n=395,393), evaluated in deciles of predicted absolute risk of ischaemic stroke.** Logistic regression was performed in the n=12,000 derivation set, adjusting for chip, sex, and 10 genetic PCs. The dashed line represents perfect calibration.

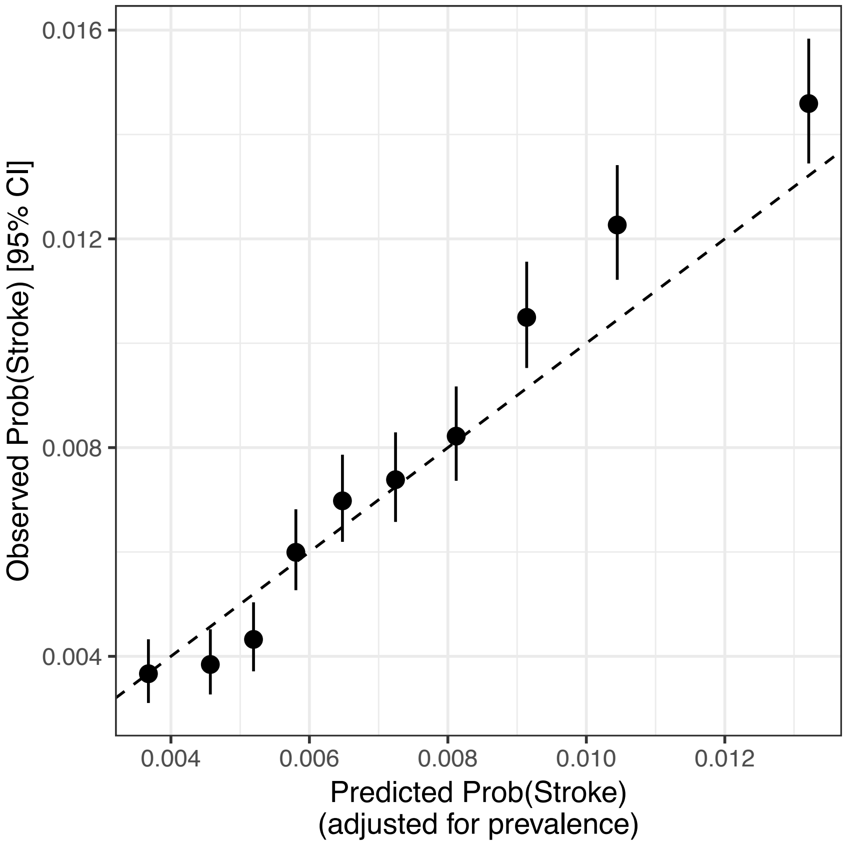

**Figure S8: The heritability explained by the metaGRS, as a function of the assumed (narrow-sense) heritability of ischaemic stroke on the liability scale.** Estimates are based on linear regression, either (i) adjusted for sex, age of assessment, and 10 genetic PCs, or (ii) not adjusted for these covariates.

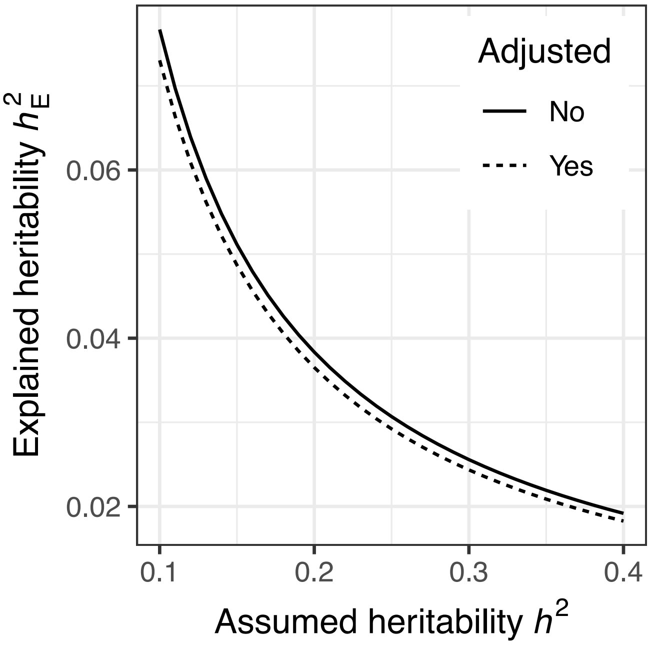

**Figure S9: Generalised additive model (GAM; with cubic splines) regression of the stroke metaGRS on the UK Biobank place of birth (north and east coordinates), n=381,041. (a)** Adjustments include: (i) unadjusted for geography; (ii); residuals from top 10 genetic PCs; (iii); residuals from top 10 PCs and natural splines of north and east coordinates; (iv); same as (iii) except adjusting for top 30 PCs; (v) residuals from 10 PCs and thin plate regression spline (TPRS) of the north and east coordinates; (vi) same as (v) but additionally adjusted for UKB assessment centre. Individuals without known place of birth were excluded. **(b)** Hazard ratios for the residuals of the metaGRS regressed on the above variables.

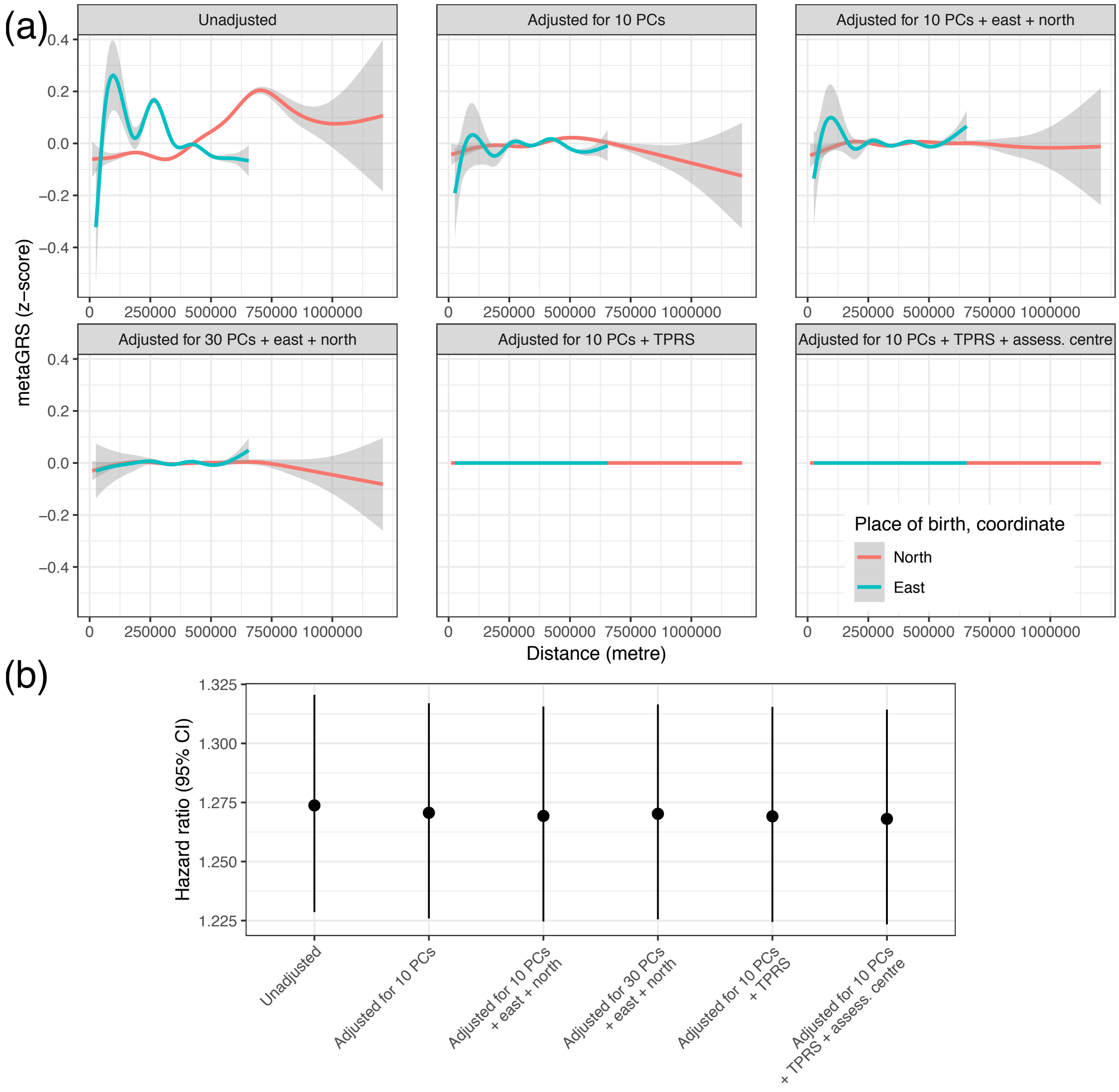

### Supplementary Tables

#### Table S1: Sources of summary statistics used for the metaGRS

| Category | GRS | Phenotype | Number of SNPs in score | Reference |
|---|---|---|---|---|
| Stroke | AS | Any stroke | 3,236,236 | Malik, et al. [4] |
| | IS | Ischaemic stroke | 3,284,492 | Malik, et al. [4] |
| | LAS | Large artery stroke | 3,353,125 | Malik, et al. [4] |
| | SVS | Small vessel stroke | 3,265,676 | Malik, et al. [4] |
| | CES | Cardioembolic stroke | 3,262,206 | Malik, et al. [4] |
| Lipids | HDL | HDL cholesterol | 115,051 | Willer, et al. [8] |
| | TG | Triglycerides | 113,406 | Willer, et al. [8] |
| | LDL | LDL cholesterol | 113,183 | Willer, et al. [8] |
| | TC | Total cholesterol | 115,019 | Willer, et al. [8] |
| Cardiovascular (non-CAD) | AF | Atrial fibrillation | 1,013 | Weng, et al. [35] |
| Smoking | Smoking | Cigarettes per day | 895,675 | Tobacco Genetics Consortium [11] |
| BP | SBP | Systolic BP | 628,674 | Wain, et al. [10] |
| | DBP | Diastolic BP | 628,601 | Wain, et al. [10] |
| Anthropometric | BMI | Body mass index | 967,600 | Locke, et al. [7] |
| | Height | Height | 965,305 | Wood, et al. [12] |
| Diabetes | T2D | Type 2 diabetes | 4,932,042 | Scott, et al. [13] |
| CAD | 46K | CAD | 45,810 | Abraham, et al. [19] |
| | FDR202 | CAD | 199 | Nikpay, et al. [44] |
| | 1KG | CAD | 1,713,315 | Inouye, et al. [17] |

**Table S2: Association of the metaGRS with established risk factors.** For continuous outcomes (SBP, BMI), the associations are from linear regression; for discrete outcomes the associations are from logistic regression. All regression models were adjusted for sex, chip, age at assessment, and 10 genetic PCs, using the validation subset of UKB (incident cases and non-cases, n=390,849).

| | **Estimate** | **95% CI** |
|---|---|---|
| Systolic BP | 1.67 | 1.61–1.73 |
| BMI | 0.44 | 0.42–0.45 |
| Family history of stroke | OR=1.06 | 1.05–1.07 |
| Diagnosed diabetes | OR=1.18 | 1.16–1.19 |
| Diagnosed high cholesterol | OR=1.21 | 1.20–1.22 |
| Current smoking | OR=1.06 | 1.05–1.07 |
| Diagnosed hypertension | OR=1.20 | 1.19–1.21 |
